# Wild gut microbiome suppresses the opportunistic pathogen *Aeromonas* in medaka under domesticated rearing conditions

**DOI:** 10.1101/2024.11.06.621463

**Authors:** Keisuke Kawano, Kai Kawabe, Yuki Sano, Tomoyuki Hori, Minoru Kihara, Yoshitomo Kikuchi, Hideomi Itoh

**Author notes:** Correspondence: Keisuke Kawano, Hideomi Itoh. Graduate School of Agriculture, Kyoto University, Sakyo-ku, Kyoto 606-8502, Japan (Keisuke Kawano). Research Center, JAPAN NUTRITION Co., Ltd., Nasu-Shiobara, Tochigi 325-0103, Japan (Yuki Sano).

## Abstract

**Background:** The gut microbiome plays a crucial role in the metabolic health and pathogen resistance of various host animals. It is also well established that external environmental factors can influence the gut microbiome, leading to differences in its composition. However, the functional implications of these differences remain poorly understood. This study examined the gut microbiome of medaka (*Oryzias latipes* species complex) by comparing domesticated and wild populations, with the aim of gaining insights into the functional significance of their specific characteristics, particularly those of the wild-type microbiome.

**Results:** For the comparative analysis of the gut microbiome, 48 domesticated and 122 wild medaka were collected from multiple laboratories, pet stores and streams across Japan. The results showed that wild medaka exhibited significantly higher gut microbiome diversity, with a broader range of bacterial members. In contrast, the gut microbiome of domesticated medaka harbored lower microbial diversity and was consistently dominated by *Aeromonas*, a typical opportunistic pathogen in fish. Additionally, 88.6% of *Aeromonas* isolates from domesticated medaka exhibited haemolytic activity. Moreover, a domesticated rearing experiment with wild populations showed no proliferation or dominance of *Aeromonas* in their gut, as observed in domesticated medaka. A further rearing experiment revealed that pre-exposing antibiotic-treated medaka to sediments from their natural habitats prevented *Aeromonas* colonisation, even when reared under domesticated conditions.

**Conclusions:** These findings suggest that the habitat-derived wild gut microbiome can inhibit *Aeromonas* proliferation in domesticated fish, highlighting its potential to mitigate opportunistic diseases in aquaculture.

## Background

Many animals host various microorganisms in their intestines, collectively known as the gut microbiome [1]. These microbial communities play a crucial role in host physiology by influencing metabolism, immune function and adaptation to environmental conditions [2–4]. For example, woodrats (*Neotoma* spp.) rely on their gut microbiota to detoxify toxic desert plants, enabling their survival in resource-limited environments [5]. Termites depend on their gut microbiota to degrade indigestible lignin and cellulose in wood, allowing them to exploit highly specialized ecological niches [6]. Similarly, the fruit fly (*Bactrocera dorsalis*) utilizes its gut microbiota to recycle urea, a nitrogenous waste, for amino acid synthesis, supporting its adaptation to nitrogen-poor habitats [7]. These examples highlight how gut microbiota perform specialized functions linked to diet and habitat, profoundly influencing host physiology and ecology.

Artificial rearing conditions, however, can induce substantial shifts in the gut microbiome due to altered diets, antibiotic use, decreased exposure to environmental microbial reservoirs and increased human interaction [8, 9]. Studies comparing animals in nature or reared in artificial environments have documented changes such as reduced microbial diversity, shifts in dominant taxa, and the loss of environmental microbial inputs [8]. However, the extent of microbiome changes varies across species. Some exhibit marked reductions in microbial diversity, leading to dysbiotic and imbalanced microbiota compared to their wild counterparts [10, 11], whereas others retain a significant portion of their wild-type microbiota, demonstrating resilience to artificial rearing conditions [12, 13]. Although some studies suggest that such alterations may compromise immune function and increase disease susceptibility [14, 15], the functional consequences of gut microbiome induced by artificial rearing conditions shifts remain unclear.

Fish gut microbiomes are similarly shaped by environmental factors such as diet, salinity, and temperature [16, 17]. Wild fish generally harbor more diverse and complex gut microbiomes, compared to domesticated fish which often exhibit reduced microbial diversity due to controlled diets, artificial rearing environments and limited exposure to environmental microbes [18–21]. The fish gut microbiota plays critical roles in development of the digestive tract, nutrient absorption, immune regulation, and disease resistance [22–24], with specific bacterial taxa contributing to metabolic and protective functions [16, 25]. For example, genera such as *Bacillus*, *Lactobacillus* and *Clostridium* are commonly found in fish gut microbiomes and suggested to be involved in nutrient cycling and pathogen defense [16, 25, 26]. Given these functions of gut microbiota and the fact that they are shaped by environmental factors, it can be inferred that different functional microbiomes are formed depending on the rearing environment and natural habitats. While studies in mammals and insects suggest that wild-derived microbiomes confer enhanced pathogen resistance [14, 27], this hypothesis has not been thoroughly investigated in fish.

In this study, we focused on medaka (*Oryzias latipes* species complex [28, 29]), a widely used experimental model and ornamental freshwater fish with readily available wild populations across Japan [30]. Domesticated medaka are typically reared in plastic or glass tanks with stable water temperatures (25°C–28°C), dechlorinated tap water, and commercially formulated artificial feed designed for rapid growth [31]. These controlled conditions contrast sharply with wild medaka habitats, which are typically muddy puddles with fluctuating temperatures and diverse food sources, including zooplankton, phytoplankton and small invertebrates [32, 33]. We performed a comparative analysis of the gut microbiota in medaka from artificial environments and natural habitats, aiming to clarify the distinct characteristics of the gut microbiota in domesticated and wild medaka, as well as explore the effect of domesticated rearing condition on wild gut microbiota and its potential role in disease risk.

## Methods

### Sample collection

We collected 48 domesticated (D) medaka from three laboratories and three pet stores (**Fig. 1**; **Table 1**). The D medaka groups included inbred/conserved lines (D1–D4 groups) and the orange– red commercial strain (himedaka, D5–D8 groups), with each group reared separately in independent tanks (**Table 1**). Three other small fish species, the guppy (*Poecilia reticulata*), goldfish (*Carassius auratus*) and zebrafish (*Danio rerio*), were obtained from the same pet store where the D8 group was purchased. All D medaka and other fish were maintained using standardised methods [31]: feeding on fishmeal-based artificial foods in dechlorinated tap water in glass or plastic tanks, with dechlorination achieved via aeration or chemical dechlorinators. Specifically, the D5 group, which is the generations of the D6 group, was maintained in our laboratory under domesticated conditions: fish were kept at 25°C in dechlorinated tap water in plastic tanks with aeration under long day conditions (14 h of light per day) and fed fishmeal-based artificial foods (TetraMin; Spectrum Brands Inc., Madison, WI, USA). Ammonium and nitrite concentrations in the rearing water were monitored daily using pack tests (Kyoritsu Chemical Lab., Tokyo, Japan); if they exceeded the safe standards for fish (ammonium, 0.2 mg/L; nitrite, 0.01 mg/L), one-third of the rearing water was replaced with fresh dechlorinated tap water. All D groups were used for the gut microbiome analysis, and each group consisted of six individuals, including three females and three males. The D8 group was also used for the isolation of *Aeromonas* strains and Rearing Experiment 2 as described below.

**Fig. 1.**
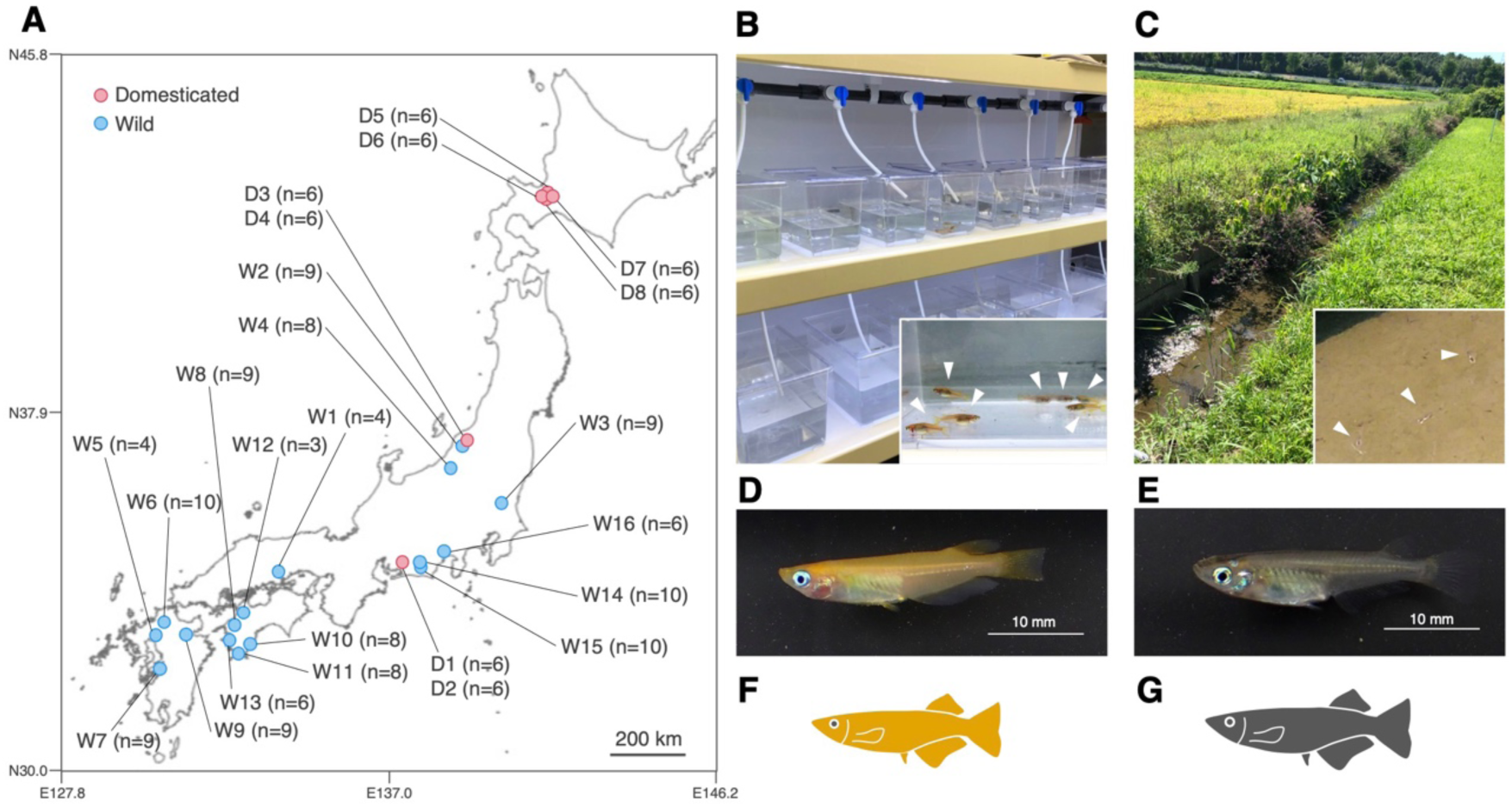
Comparison of habitat and morphology between domesticated (D) and wild (W) groups of *Oryzias latipes* species complex in Japan. (A) Geographical locations of collection sites for D and W groups. Numbers in parentheses indicate the number of individuals used in the gut microbiome analysis, as shown in Fig. 2. Detailed information is summarized in **Table 1**. (B) Typical laboratory rearing conditions of the D groups. The inset photo shows medaka from the D groups (closed triangles). (C) Stream habitat of W groups. The inset photo showing wild medaka in a mud puddle (closed triangles). Photographs of the fish specimen in the D8 (D) and the W7 groups (E). Icons representing D medaka (F) and W medaka (G) used in subsequent figures. Map data sourced from the Geospatial Information Authority of Japan.

**Table 1.**
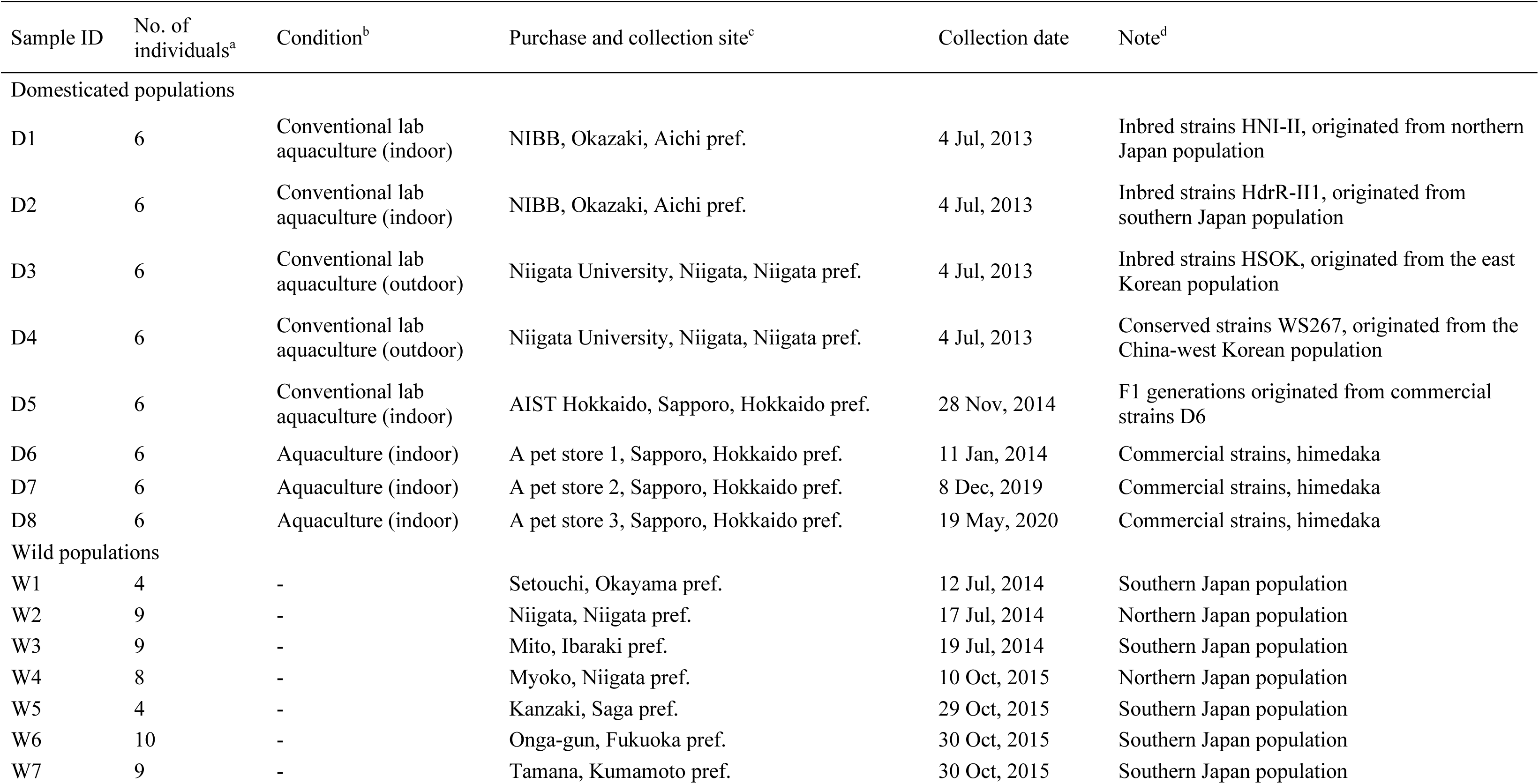
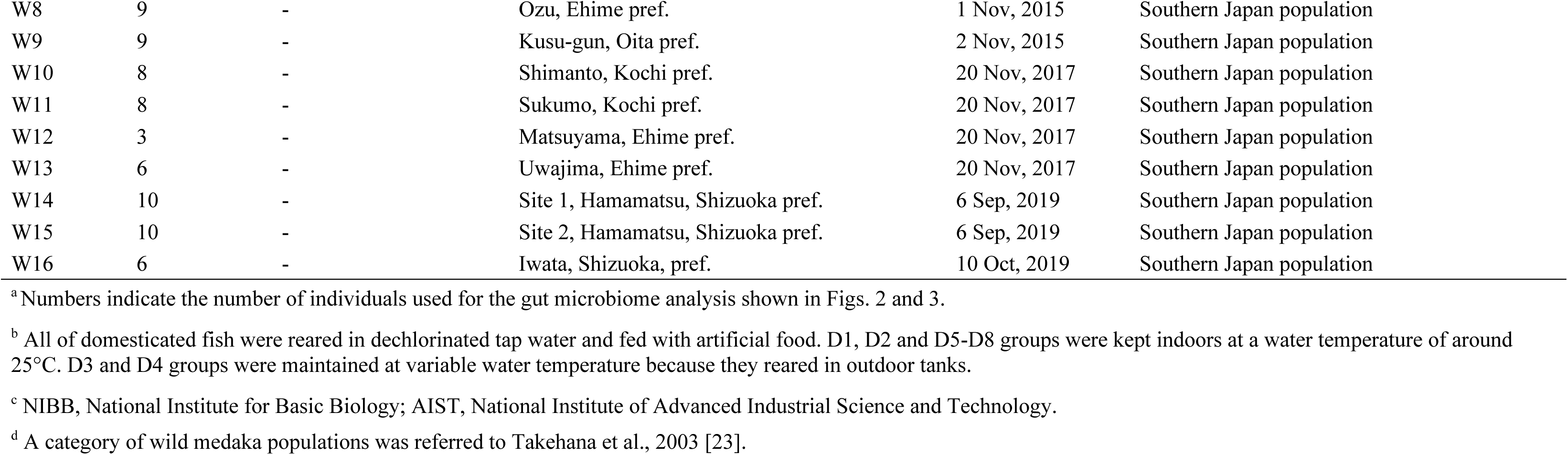
Medaka examined in this study.

The W medaka groups (W1–W16), consisting of 122 individuals, were caught using nets in 16 streams across 10 prefectures in Japan from summer to fall between 2014 and 2019 (**Fig. 1**; **Table 1**). The collected individuals were transported alive to the laboratory within two days and were used for dissection to analyse the gut microbiome (W1–W16 groups) or for the subsequent rearing experiment (W10–W13 groups, Rearing experiment 1 as described below). Stream sediments (S6, S12, S14 and S15) were also collected from the sites where the W6, W12, W14 and W15 medaka groups were captured. The mud from the stream was scooped up with a shovel, and water-saturated sediment from a depth of 0-5 cm was collected. The sediment was then transported to the laboratory and stored at 4°C until used for Rearing Experiment 2 as described below.

All fish in D and W groups analysed were adults over 2 cm in length. Comprehensive approval for the experimental use of fish was granted by the institutional animal care and use committee. Sampling and experiments followed the ‘Guidelines for the Use of Fishes in Research’ established by the Ichthyological Society of Japan in 2003 (http://www.fish-isj.jp/english/guidelines.html).

### Rearing Experiment 1: Domesticated rearing of wild medaka

After the W10–W13 groups were transported alive from field sites to the laboratory within two days, a subset of medaka from each group (n = 6 per group) was kept separately in a 10-L plastic tank (15.3 × 27.8 × 16.5 cm) for 30 days under the same domesticated conditions as the D5 group, as described above. Specifically, the tanks were maintained at 25°C with 8 L of dechlorinated tap water under long day conditions and the individuals were fed 0.5 g of TetraMin flakes (Spectrum Brands) per tank daily. Ammonium and nitrite concentrations in the rearing water were monitored and managed as described above. After 30 days, the domestically reared wild medaka groups (DW10–DW13) were dissected for gut microbiome analysis, as described below.

### Rearing Experiment 2: Domesticated rearing of medaka with transplanted environmental microbes

#### Step 1: Antibiotic treatment of medaka

Two individuals from the D8 group were placed in pairs in a 250-mL sterile plastic flask with a vented cap (ASONE, Osaka, Japan) containing 200 mL of sterile gnotobiotic zebrafish medium (GZM) [34] with a 1% penicillin/streptomycin/amphotericin B mixture (FUJIFILM Wako Pure Chemical Corporation) and 100 µL/mg erythromycin (Tokyo Chemical Industry Co., Tokyo, Japan) for 24 h. After treatment, each medaka was rinsed three times in 200 mL of fresh sterile GZM for 10 min to remove residual antibiotics. The treated medaka were then used in the next step, i.e., for ‘Step 2: Pre-rearing with stream sediments’ as described below. Sterility was confirmed using a culture-based method for germ-free zebrafish [34]. The guts were dissected, homogenised in 100 µL of sterile GZM, diluted and spread on 1.5% agar plates containing Luria– Bertani broth (Becton, Dickinson and Company, Sparks, MD, USA), Nutrient Broth (Becton, Dickinson and Company) and R2A Broth (Nihon Pharmaceutical Co., Tokyo, Japan). After incubation at 25°C for 3 days, colony-forming units (CFUs) were measured.

#### Step 2: Pre-rearing with stream sediments

Initially, 200 g of each sediment (S6, S12, S14 and S15), consisting of water-saturated mud collected from streams where wild medaka (W6, W12, W14 and W15 groups) were caught (**Table 1**), was added to 6 L of sterile dechlorinated tap water in separate 10-L plastic tanks (i.e., one tank for each sediment type). After aerating the tanks at 25°C for one week, the antibiotic-treated medaka, prepared as described above, were reared in these tanks for one week. Following this period, the medaka gut, sediment, and rearing water were used for microbiome analyses. As a control, the antibiotic-treated medaka were reared in 6 L of sterile dechlorinated tap water without sediment for one week, with two separate tanks prepared for this condition.

#### Step 3: Domesticated rearing

Medaka pre-reared with each sediment (DS6, DS12, DS14 and DS15 groups) or without sediment (C1 and C2 groups) were rinsed in 200 mL of fresh sterile dechlorinated tap water three times for 10 min to remove residual sediment, and each group was then transferred to a dedicated 17-L plastic tank (20.8 × 31.4 × 27.0 cm) with 10 L of rearing water routinely used for maintaining the D5 group. The microbiome of this rearing water was also analysed. The medaka were reared for three weeks under the same domesticated conditions as the D5 group, as described above, and were then used for gut microbiome analysis.

### DNA extraction

All fish were fasted for 1–3 days to remove transient gut contents and then anesthetised on ice. After rinsing twice with ice-cold sterile distilled water, the entire gut was dissected using ethanol-treated forceps and tweezers under a binocular microscope, transferred to a microcentrifuge tube and homogenised with 180 µL of AL Buffer (Qiagen, Hilden, Germany) using a pestle. After 1 h at 37°C with 20 µL of 20 mg/mL lysozyme (FUJIFILM Wako Pure Chemical Corporation, Osaka, Japan), the homogenates were incubated overnight at 56°C with 20 µL of 20 mg/mL protease K (Qiagen). The crude extract was purified using the QIAamp DNA Mini Kit (Qiagen). Water samples (50–100 mL of rearing water) were first filtered through a 10 µm pore size membrane to remove suspended debris, and then through a 0.22 µm pore size membrane to capture bacterial cells. DNA extraction from the resulting membranes and sediment samples followed the same method as for gut DNA extraction. All DNA samples were stored at −30°C until use.

### Diversity analysis of the microbiome

The variable V4 region of the bacterial 16S rRNA gene was amplified using the universal primer set 515F with Illumina P5 sequences and 806R with 12-base indexes and Illumina P7 sequences [35], Q5 Hot Start High-Fidelity Master Mix (New England Biolabs, Beverly, MA, USA) and the prepared DNA, as described previously [36]. Following 1.5% agarose gel electrophoresis using Novel Juice dye (GeneDireX, Las Vegas City, NV, USA), appropriately sized PCR amplicons were cut from the gels and purified using the Wizard SV Gel and PCR Clean-Up System (Promega, Madison, WI, USA). Each purified DNA sample was quantified using a Qubit 2.0 Fluorometer (Invitrogen, Carlsbad, CA, USA), diluted and mixed equally. Paired-end sequencing of the PCR amplicon libraries was conducted with a PhiX control on a MiSeq or iSeq sequencing platform (Illumina) using the MiSeq Reagent Kit v2 or iSeq 100 Reagent Kit (both Illumina), respectively, following the manufacturer’s instructions. Raw sequence preprocessing and quality filtering, including paired-end sequence joining and removal of low-quality and chimeric sequences, were performed as previously described [37]. Taxonomic assignment was performed using the RDP classifier ver. 2.13 [38] with a 50% confidence threshold.

### Quantitative analysis of the microbiome

Quantitative PCR (qPCR) was performed to amplify the V4 region of the bacterial 16S rRNA gene using the universal primers 515F and 806R, along with a LightCycler 96 System (Roche Diagnostics, Basel, Switzerland) and THUNDERBIRD SYBR qPCR Mix (Toyobo, Otsu, Shiga, Japan), following the manufacturers’ instructions. The thermal cycling conditions were as follows: initial denaturation at 95°C for 30 s, followed by 45 cycles at 95°C for 10 s, 60°C for 30 s and 72°C for 30 s. The gene copy number of the bacterial 16S rRNA gene was calculated from a standard curve based on a dilution series of *Aeromonas hydrophila* NBRC13286 PCR products. Additionally, the abundance of *Aeromonas* was calculated by multiplying the bacterial abundance through this qPCR by the proportion of *Aeromonas* in gut microbiome obtained from the diversity analysis described above.

### Isolation and haemolysis assay of *Aeromonas* strains from the medaka gut

To examine the pathogenic potential of *Aeromonas* in the D medaka gut, we isolated *Aeromonas* strains and subjected them to a haemolysis assay on blood agar. Whole guts were dissected from D8 samples and homogenised in 100 µL of GZM using a pestle. The homogenates were diluted with GZM and spread on Trypticase soy agar containing 5% sheep blood (Becton, Dickinson and Company). After incubation at 25°C for 3 days, halo formation around the colonies was monitored for haemolysis activity. Colonies were picked and subjected to 16S rRNA gene sequencing, as described previously [39]. Haemolytic *Aeromonas* strains were cultured overnight in Luria– Bertani medium at 27°C and washed with GZM. After adjusting the concentration to OD_600_ = 0.01 using GZM, 10 µL of the bacterial suspension was dropped onto Trypticase soy agar with 5% sheep blood (Becton, Dickinson and Company) and cultured at 27°C for 3 days. Haemolysis activity was evaluated based on the diameter of the halo formed around the drop. Halo size was measured using Fiji [40]. *Aeromonas hydrophilia* NBRC 13286 was used as the control in the haemolysis assay.

### Statistical analysis

Beta and alpha diversity were analysed by normalising read counts to match the sample with the minimum read counts using MicrobiomeAnalyst 2.0 [41]. Beta diversity at the bacterial genus level among the gut microbiota was assessed using the Bray–Curtis dissimilarity index. Nonmetric multidimensional scaling (NMDS) was used for data visualization. Differences in beta diversity were statistically evaluated using PERMANOVA with sample type (D vs W, W vs DW, D5 vs DW or C vs DS groups), using 999 permutations to assess statistical significance. Alpha diversity was calculated using the Chao1 and Shannon indices. To identify specific bacterial groups in medaka under each condition, linear discriminant effect size (LEfSe) analysis was conducted using MicrobiomeAnalyst 2.0 [41]. For pairwise comparisons, the alpha indices and *Aeromonas* proportions/abundance in the gut between each pair of groups, as well as CFUs before and after antibiotic treatment, were analysed using the Mann–Whitney U-test (package “exactRankTests” in R). For comparisons across multiple group, the haemolytic activity of *Aeromonas* strains was analysed using the Mann–Whitney U-test with Bonferroni correction. These statistical comparisons were conducted using R studio 4.2.2. [42].

### Availability of nucleotide sequence data

Illumina sequencing data from all medaka gut, sediment and water samples were available in the DDBJ/GenBank/EBI databases under accession PRJNA1102999. Additionally, Sanger sequencing data from isolated *Aeromonas* strains were available in the same databases under accessions PP474260, PP474261 and PP549924–PP549964.

## Results

### Domesticated and wild medaka gut microbiome profiles

The NMDS plot based on the Bray–Curtis distance revealed a clear distinction in gut microbiome community structures between domesticated and wild populations (PERMANOVA: *F* = 42.043, *r*^2^ = 0.200, *p* < 0.05; **Fig. 2A**). Alpha diversity indices, Chao1 and Shanonn indices, showed that the D groups had lower microbiome richness and evenness compared with the W groups (**Fig. 2B,C**), indicating greater diversity in wild populations. Taxonomic analysis at the phylum level revealed *Proteobacteria* as the dominant group in both D and W groups (**Fig. 2D**), whereas *Fusobacteria* were also abundant in half of the D groups (D5–D8) (**Fig. 2D**). Depending on the collection site, the W groups exhibited a more diverse microbiome, including phyla such as *Planctomycetes*, *Verrucomicrobia*, *Firmicutes*, *Chloroflexi*, *Acidobacteria* and *Actinobacteria*, which were less prominent in the D groups (**Fig. 2D**).

**Fig. 2.**
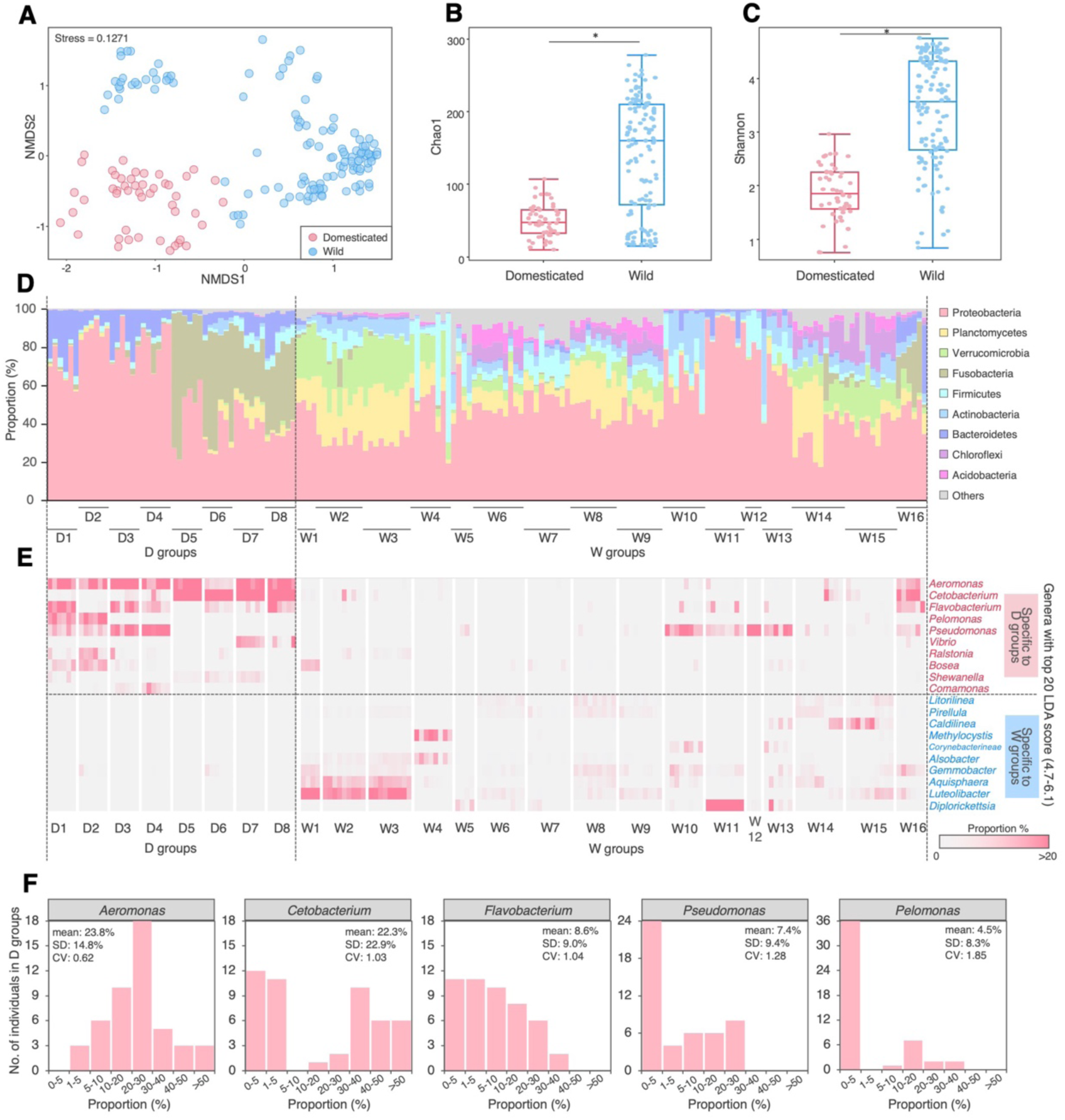
Diversity and specificity of the gut microbiome of medaka between the D and W groups. (A) Nonmetric multidimensional scaling (NMDS) plot illustrating gut microbiome dissimilarity based on the Bray–Curtis distance. Microbial diversity comparison using Chao1 (B) and Shannon (C) indices. Asterisks indicate significant differences (*p* < 0.05, Mann–Whitney U-test). (D) Community structure of the gut microbiome according to bacterial phylum proportions. (E) Specific bacterial groups associated with the D or W groups, identified by LEfSe analysis, listing the top 10 genera with the highest LDA scores. (F) Histograms showing the top 5 groups in the D groups. SD, standard deviation; CV, coefficient of variation.

LEfSe analysis identified genera specific to each group. *Aeromonas*, *Cetobacterium* and *Flavobacterium* were specific to the D groups, whereas *Diplorickettsia*, *Luteolibacter* and *Aquishaera* were unique to W groups (**Fig. 2E; Supplementary Fig. S1**). The proportion of *Cetobacterium* was high (>30%) in most D5–D8 groups (91.7%, 22/24 individuals), but lower (<5%) in most D1–D4 groups (95.8%, 23/24 individuals) (**Fig. 2E,F**). In the W groups, *Diplorickettsia* was predominantly found in the W11 group (33.7%–83.1%, 8 individuals) but rarely in other W groups (0%–8.4%, 116 individuals) (**Fig. 2E**). *Aeromonas*, which had the highest linear discriminant analysis (LDA) score and average proportion (**Supplementary Fig. S1A,B**), was consistently detected in the D groups (**Fig. 2E**; **Supplementary Fig. S1B**), confirming it as the most common genus in domesticated medaka. This is further supported by its lowest coefficient of variation (CV) among the dominant genera in the D groups (**Fig. 2F**), indicating its relatively stable abundance.

### Potentially pathogenic *Aeromonas* in the gut of domesticated fish

*Aeromonas* was the dominant bacterial member in the gut of D groups (**Fig. 2E,F; Supplementary Fig. S1B**), accounting for 4.2%–80.4% of the gut microbiome (mean ± SD = 23.8% ± 14.8%, n = 48; **Fig. 3A**). However, *Aeromonas* levels in the W groups were consistently low (0%–8.6%; mean ± SD = 0.5% ± 1.2%, n = 116), except for the W16 group where they were slightly higher (1.9%–15.8%; mean ± SD = 9.1% ± 4.5%, n = 6; **Fig. 3A**). Overall, *Aeromonas* levels were significantly higher in the D groups compared with the W groups (**Fig. 3B**). Quantitively, *Aeromonas* levels in the D groups ranged from 1.3 × 10^6^ to 1.3 × 10^9^ copies/individual (mean ± SD = 2.0 ± 2.8 × 10^8^ copies/individual, n = 48), whereas in the W groups they were substantially lower (3.0 × 10^7^ to 1.2 × 10^7^; mean ± SD = 0.4 ± 1.5 × 10^6^ copies/individual, n = 116) (**Fig. 3C**). Both the proportion and quantity of *Aeromonas* were significantly higher in the gut of D groups (**Fig. 3D**).

**Fig. 3.**
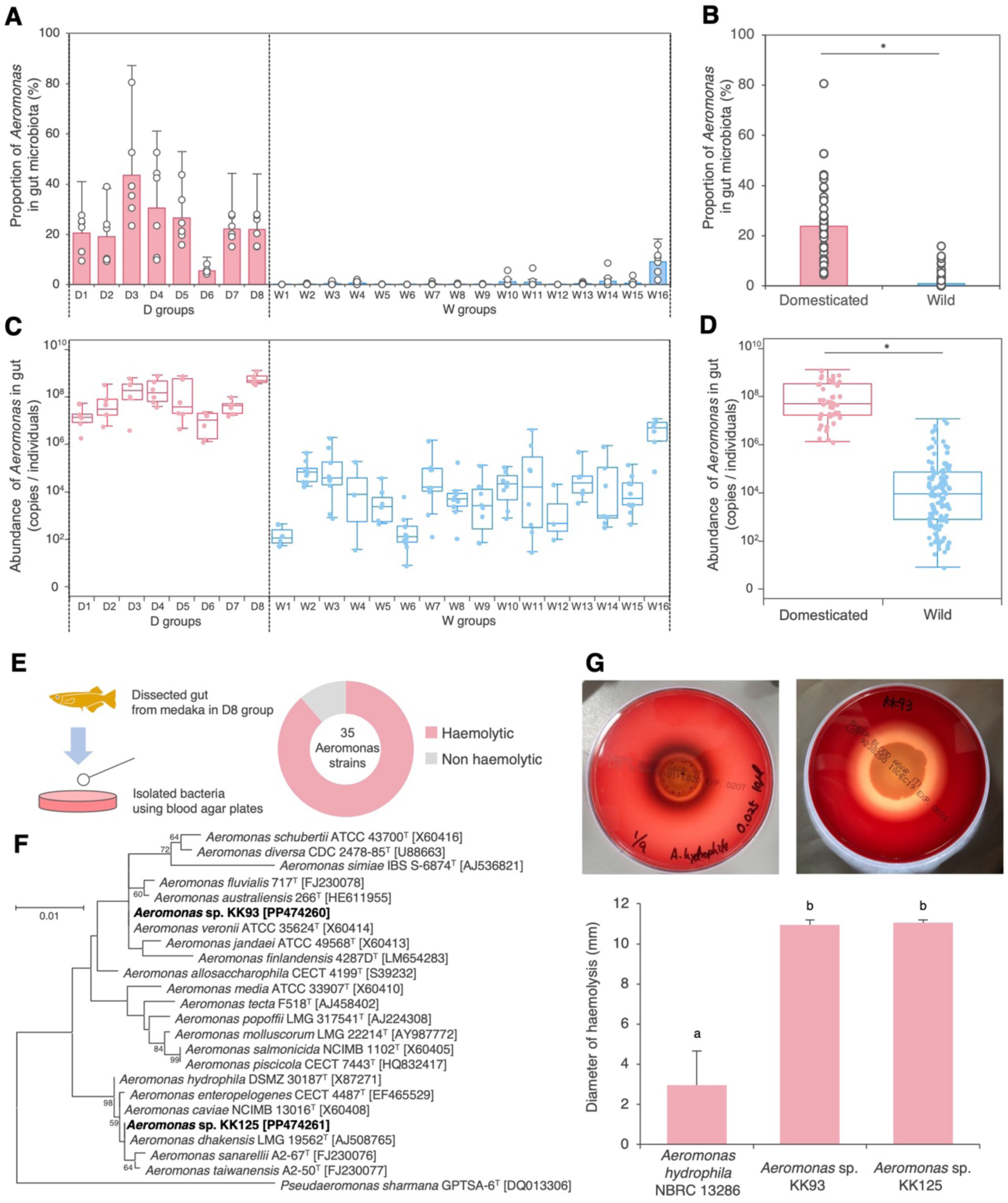
Distribution and potential toxicity of *Aeromonas* in the medaka gut. (A) Proportion of *Aeromonas* in the gut microbiome of medaka samples in each group. (B) Average proportion of *Aeromonas* compared between the D and W groups. (C) Abundance of *Aeromonas* in the gut microbiome, estimated via amplicon sequencing and qPCR analyses. (D) Average abundance of *Aeromonas* compared between D and W medaka. The asterisks in (C) and (D) indicate statistically significant differences (*p* < 0.05, Mann–Whitney U-test). (E) Schematic overview of haemolytic bacterial strain isolation from the medaka gut in the D8 group, highlighting the composition of 35 *Aeromonas* strains. Isolates other than *Aeromonas* are summarised in **Supplementary table S1**. (F) Phylogenetic tree of *Aeromonas* strains isolated from the guts of domesticated medaka, based on 16S rRNA gene sequences. *Pseudaeromonas sharmana* GPTSA-6^T^ was used as the outgroup. Scale bar represents 0.01 substitutions per nucleotide position. (G) Haemolytic activity of *A. hydrophila* and medaka-derived *Aeromonas* strains, assessed via halo formation on blood agar (right photo: *A. hydrophila* NBRC 13286; left photo: *Aeromonas* sp. strain KK93). A bar graph displays the mean with standard deviation from six replicates, with different letters indicating statistically significant differences (*p* < 0.05, Mann–Whitney U-test with Bonferroni correction).

Additionally, we analysed guppy (*Poecilia* sp.), goldfish (*Carassius* sp.) and zebrafish (*Danio rerio*) from the pet store as the D8 group. *Aeromonas* proportions in the gut microbiome were 5.0%–10.9% in guppy (mean ± SD = 7.5% ± 2.2%, n = 6), 10.4%–92.3% in goldfish (mean ± SD = 49.6% ± 20.6 %, n = 6) and 15.0%–27.9% in zebrafish (mean ± SD = 17.0% ± 5.8%, n = 6) (**Supplementary Fig. S2**).

Specific *Aeromonas* species are opportunistic pathogens that cause diseases, such as haemorrhagic inflammation, ulcers and fin rot in fish [43]. Of the 35 *Aeromonas* strains isolated from the D5 group (**Fig. 3E; Supplementary Table S1**), 88.6% (31/35 strains) showed haemolytic activity, forming halos on the blood agar (**Fig. 3E**). Among 40 haemolytic isolates, 77.5% (31/40 strains) were *Aeromonas* (**Supplementary Table S1**). Two representative *Aeromonas* isolated strains, *A. veronii* KK93 and *A. dhakensis* KK125 (**Fig. 3F; Supplementary Table S1**), exhibited stronger haemolytic activity compared with *A. hydrophila* NBRC13286 isolated from the intestine of juvenile silver salmon (**Fig. 3G**).

### Rearing Experiment 1: Gut microbiome of wild medaka reared under domesticated conditions

Rearing wild medaka in each W10–W13 group under the same domesticated conditions as D5 group (generating ‘domesticated wild medaka’, DW10–DW13 groups; **Fig. 4A**) resulted in significant shifts in their gut microbiome composition. An NMDS plot based on Bray–Curtis distances revealed that the gut microbiome of DW groups differed significantly from that of both relevant W and D5 groups (PERMANOVA: DW vs. W: *F* = 31.01, *r*^2^ = 0.41342, *p* < 0.05; DW vs. D5: *F* = 25.626, *r*^2^ = 0.50618, *p* < 0.05; **Fig. 4B**). The predominant gut bacteria also shifted markedly after domesticated rearing: although *Pseudomonas*, *Acinetobacter* and *Diplorickettsia* were dominant in the W10–W13 groups, these genera were nearly absent in the DW10–DW13 groups, replaced by *Aquisphaera*, *Luteolibacter* and *Legionella* among others (**Fig. 4C**). The specificity of these genera to W or DW groups was supported through LEfSe analysis (**Fig. 4D**).

**Fig. 4.**
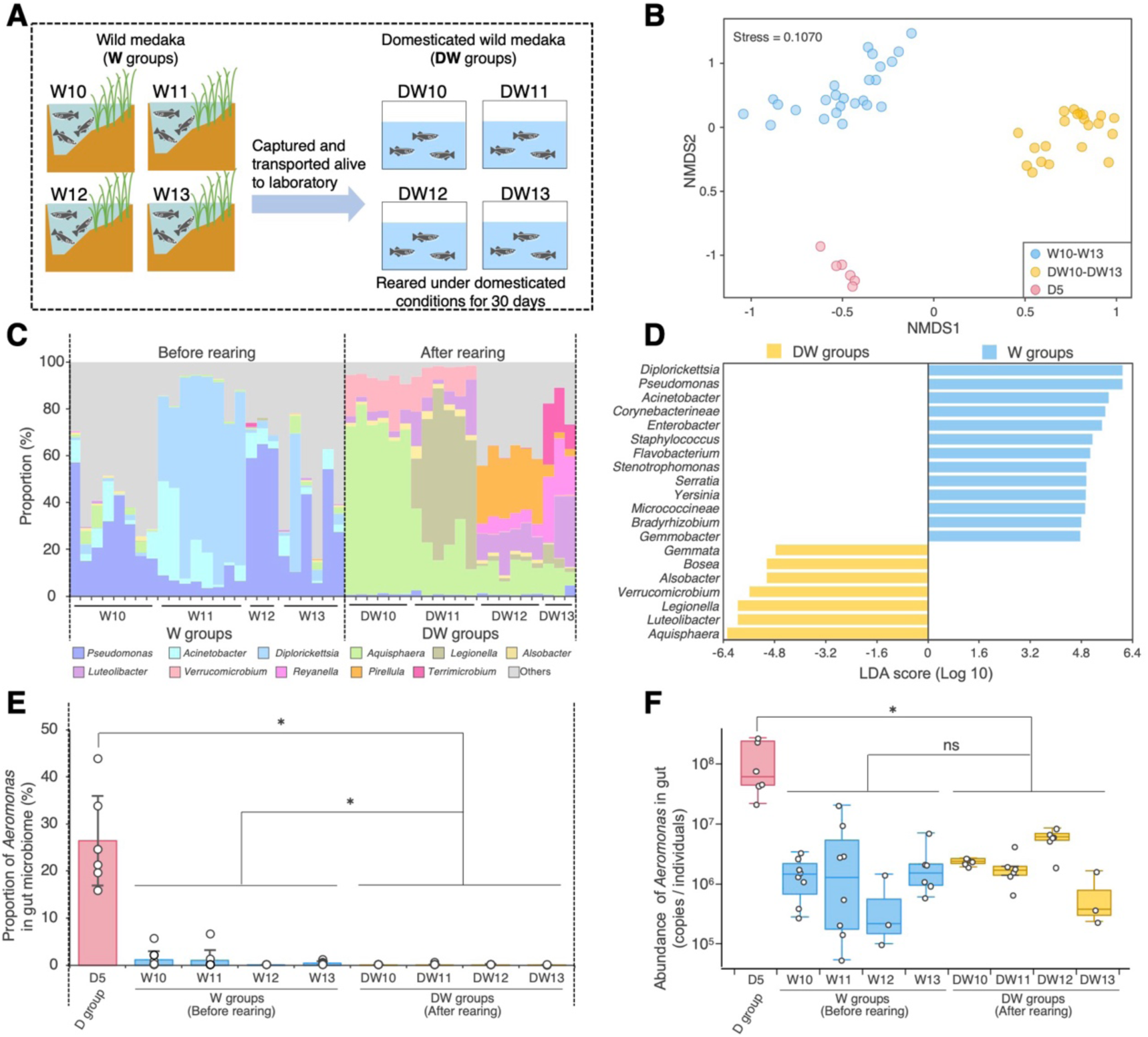
Gut microbiome of wild medaka reared under domesticated conditions. (A) Schematic overview of Rearing Experiment 1: the experimental design for preparing domestically reared wild (DW) medaka. The W10–W13 groups, collected from four different locations (Table 1), were placed in separate tanks and reared under domesticated conditions for 30 days. (B) NMDS plot based on the Bray–Curtis distance, illustrating dissimilarity in the community structure of the gut microbiome. (C) Community structure of the gut microbiome at the bacterial genus level before (W10–W13) and after (DW10–DW13) rearing under domesticated conditions. (D) Specific bacterial groups in W or DW medaka identified via LEfSe analysis. Genera with an LDA score >4.0 are shown. (E) Proportion of *Aeromonas* in the gut microbiome of each medaka sample. (F) *Aeromonas* abundance in the gut microbiome of each medaka sample, estimated through amplicon sequencing and qPCR analyses. The asterisks in (E) and (F) indicate statistically significant differences (*p* < 0.05, Mann–Whitney U-test). Data for D5 and W10–W13 groups are the same as in Fig. 2.

Notably, although the DW groups were reared under the same domesticated conditions as the D5 group, the proportion of *Aeromonas* in their gut of the DW groups (mean ± SD = 0.1% ± 0.1%, n = 21) remained significantly lower than in the D5 group (mean ± SD = 26.5% ± 9.5%, n = 6) and was even lower than in the corresponding W groups (mean ± SD = 0.8% ± 1.6%, n = 25) (**Fig. 4E**). While the quantity of *Aeromonas* in the DW groups (mean ± SD = 1.4 ± 1.9 × 10^5^, n = 21) did not differ from that in the corresponding W groups (mean ± SD = 2.6 ± 8.6 × 10^5^, n = 25), it remained lower than in the D5 group (mean ± SD = 2.4 ± 3.1 × 10^8^, n = 6) (**Fig. 4F**).

### Rearing Experiment 2: Effect of sediment pretreatment on the gut microbiome of medaka reared under domesticated conditions

Based on the results of Rearing Experiment 1, we conducted an additional experiment to examine the influence of environmental microbes on the dominance of *Aeromonas* in the gut by pre-exposing medaka to these microbes before domesticated rearing (**Fig. 5A**).

**Fig. 5.**
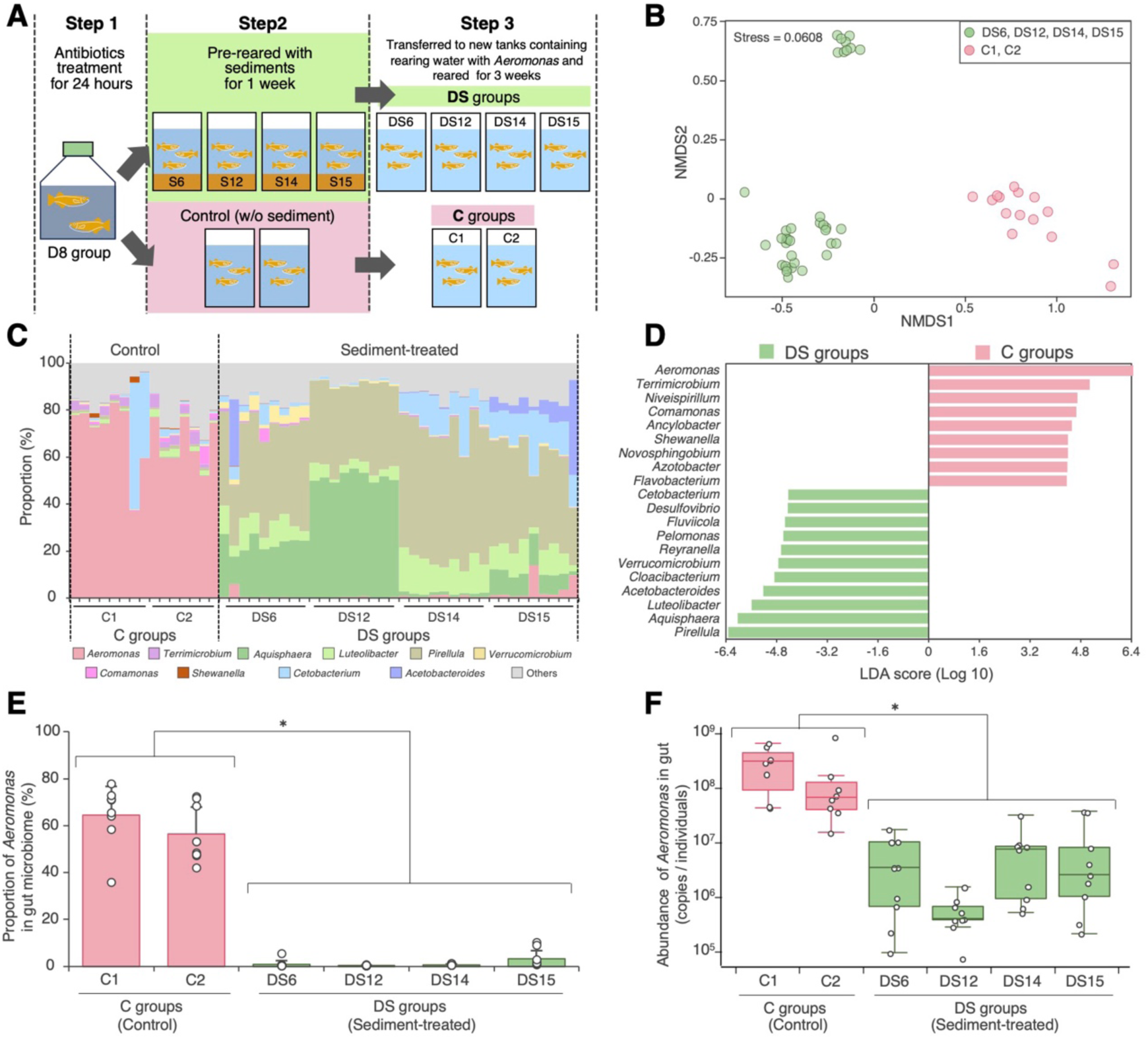
Effect of sediment pretreatment on the gut microbiome of medaka reared under domesticated conditions. (A) Schematic overview of Rearing Experiment 2: the experimental design for DS groups (DS6, DS12, D14 and D15), i.e., domesticated medaka (D8 group) pre-reared with sediment from W6, W12, W14 and W15 medaka habitats. (B) NMDS plot illustrating gut microbiome dissimilarity based on the Bray– Curtis distance. (C) Community structure of the gut microbiome at the bacterial genus level with (DS groups) or without (C groups) sediment pretreatment. (D) Specific bacterial groups in the DS or C groups identified via LEfSe analysis. Genera with an LDA score >4.0 are shown. (E) Proportion of *Aeromonas* in the gut microbiome of each medaka sample. (F) *Aeromonas* abundance in the gut microbiome of each medaka sample, estimated through amplicon sequencing and qPCR analyses. The asterisks in (E) and (F) indicate statistically significant differences (*p* < 0.05, Mann–Whitney U-test).

Prior to rearing with field-derived environmental microbes, medaka in the D8 group underwent antibiotic treatment to remove their existing gut microbiota (**Step 1 in Fig. 5A**). This reduced CFUs on three types of agar media to less than 1/10,000 of the initial CFUs (**Supplementary Fig S3**), with no adverse effects on medaka health. The antibiotic-treated medaka were then pre-reared for one week with each sediment (S6, S12, S14 and S15) from the streams where wild medaka (W6, W12, W14 and W15) were captured (**Step 2 in Fig. 5A**). After this period, genera such as *Aquisphaera*, *Zavarzinella*, *Pirellula* and *Reyranella*, originating from the sediments, colonised the gut as well as rearing water (**Supplementary Tables S2–S5**).

These sediment-treated medaka (DS6, DS12, DS14 and DS15 groups) were transferred into new tanks filled with rearing water containing *Aeromonas* (**Supplementary Table S6**) and reared for an additional three weeks under domesticated conditions (**Step 3 in Fig. 5A**). After rearing, the gut microbiome in the DS groups structure differed significantly from that of control medaka (C1 and C2 groups) reared without sediment exposure (PERMANOVA: DS vs. C: *F* = 100.71, *r*^2^ = 0.6727, *p* < 0.05; **Fig. 5B**). The duplicate control groups exhibited gut microbiomes dominated by *Aeromonas* (C1: mean ± SD = 64.5% ± 12.2%, n = 8; C2: mean ± SD = 57.7% ± 11.8%, n = 7), similar levels to the D groups (**Fig. 3A,B**), with *Aeromonas* being the most dominant genus in almost all control samples (14/15 individuals; **Fig. 5C**). In contrast, the gut microbiomes of the DS groups were dominated by *Pirellula*, *Aquisphaera* and/or *Luteolibacter* (**Fig. 5C**). The specificity of these predominant genera in C or DS groups was confirmed statistically via LEfSe analysis (**Fig. 5D**). The proportion of *Aeromonas* in the gut of DS medaka (mean ± SD = 1.3% ± 2.2%, n = 36) was significantly lower than that in the C groups (mean ± SD = 61.3% ± 12.5%, n = 15) (**Fig. 5E**). These levels were comparable to those found in the W groups (**Fig. 3A,B**). Additionally, the quantity (copies/individual) of *Aeromonas* in the DS medaka gut (mean ± SD = 6.1 ± 9.9 × 10^6^, n = 36) was two orders of magnitude lower than that in the controls (mean ± SD = 2.6 ± 2.6 × 10^8^, n = 15) (**Fig. 5F**).

## Discussion

This study reports the difference in gut microbiome diversity between domesticated and wild medaka (**Fig. 1**). Wild medaka had a significantly higher microbial diversity than domesticated medaka (**Fig. 2B,C**), with a broader range of bacterial phyla, including *Planctomycetes*, *Verrucomicrobia*, *Firmicutes* and *Acidobacteria*, which were largely absent in domesticated medaka (**Fig. 2D**). In contrast, domesticated medaka exhibited reduced diversity and a higher abundance of Proteobacteria and Fusobacteria, along with consistent *Aeromonas* dominance (**Fig. 2E,F; Supplementary Fig. S1A,B**). This trend aligns with previous studies showing higher gut microbiome diversity in wild versus domesticated animals [44–47], with the pattern likely due to more varied diets and exposure to diverse environmental microbes.

The dominance of *Aeromonas* in the gut of domesticated medaka has also been reported in previous studies [48, 49]. This trend is commonly observed in other domesticated freshwater fish species, including zebrafish, guppy, rainbow trout and various cyprinid species [26, 44, 50–54] (**Supplementary Fig. S2**). However, in wild fish, such as guppy, stickleback, gizzard shad and cyprinid species, *Aeromonas* is less dominant [44, 51, 55–57]. These findings suggest that the reduced dominance of *Aeromonas* in wild populations, compared to their domesticated counterparts, may be relatively common across freshwater fish species.

Several factors may explain *Aeromonas* proliferation in the guts of domesticated fish. First, *Aeromonas* is a mesophilic bacterium, with the rearing temperature for medaka (24°C– 28°C) providing optimal conditions for its growth [58]. Additionally, the provision of nutrient-rich artificial feed supports its growth, with *Aeromonas* known to thrive under nutrient-abundant conditions [58]. Furthermore, overcrowded conditions in domesticated rearing environments may weaken immune defences, making it easier for opportunistic pathogens, including *Aeromonas*, to proliferate. Although these factors likely encourage *Aeromonas* dominance in domesticated fish, with this phenomenon observed in various fish species, the present study revealed that wild medaka, even when reared under domesticated conditions, did not exhibit *Aeromonas* proliferation or dominance in the gut (Rearing Experiment 1, **Fig. 4**). This suggests that the wild gut microbiome may exert stronger selective pressure on *Aeromonas* relative to domesticated rearing conditions. Longer-term studies could clarify how sustained *Aeromonas* suppression by the wild gut microbiome occurs under domesticated conditions.

A notable finding is that introducing environmental microbes into medaka successfully suppressed *Aeromonas* dominance under domesticated rearing conditions (Rearing Experiment 2, **Fig. 5**). In the D8 groups, which typically exhibit *Aeromonas* dominance under these conditions (**Figs. 2E and 3A**), pretreatment with sediment from wild medaka habitats prevented *Aeromonas* from becoming dominant (**Fig. 5**). This suggests that the established gut microbiota has a stronger influence on *Aeromonas* suppression than either the medaka’s genetic background or rearing environment. This suppression can be explained by the priority effect, a phenomenon in which established microbial communities prevent the invasion of new microbes [59]. This effect has been widely observed in the guts of various animals and in plant rhizospheres [60]. In the present study, the microbes introduced from the sediment may have outcompeted *Aeromonas* for resources, occupied key ecological niches, or modified the gut environment in a way that hindered *Aeromonas* colonisation. Therefore, this study emphasises the critical role of the environmental microbiome in preventing *Aeromonas* overgrowth, highlighting the importance of the priority effect.

*Pirellula*, *Aquisphaera* and *Luteolibacter* were consistently associated with wild medaka, domesticated wild-reared medaka and sediment-exposed medaka (**Figs. 2, 4 and 5**), suggesting that they may inhibit *Aeromonas* through priority effects and/or direct antagonistic interactions. *Pirellula* and *Aquisphaera* belong to the phylum Planctomycetes, commonly found in aquatic and terrestrial environments [61, 62], and are frequently detected in fish gut microbiomes in natural or semi-natural settings [63, 64]. *Luteolibacter*, a Verrucomicrobia member, is found in diverse environments, including seawater, soil and activated sludge [65–67], as well as the guts of annelids, including the medicinal leech [68]. Given their association with lower *Aeromonas* levels, these bacteria may use antibiotics or injectors (such as those from the Type VI secretion system) to attack *Aeromonas*, or they may form robust communities, such as biofilms, that prevent its colonisation. However, the ecological roles of these bacteria are not well understood; therefore, future research should isolate strains from medaka guts and test their antagonism against *Aeromonas* through coculture experiments *in vitro* and i*n vivo*.

The consistent dominance of *Aeromonas* in domesticated medaka is an important finding that should also be highlighted from the perspective of applied science. Although *Aeromonas* can aid digestion and immune activation [69], it is more commonly recognised as an opportunistic pathogen in fish [70, 71]. Environmental stressors, such as water quality or temperature changes, can trigger *Aeromonas*-related infections [72], with a high abundance of *Aeromonas* in the gut increasing infection risk. In this study, 88.6% of *Aeromonas* isolates from domesticated medaka exhibited haemolytic activity (**Fig. 3E**), suggesting a potential health risk for these fish. In aquaculture, efforts are being made to reduce the prevalence of *Aeromonas* in fish and their environments given the pathogen’s role in disease outbreaks [70, 71, 73]. Current *Aeromonas* control methods rely on antibiotics, but this bacterium is highly prone to developing antibiotic resistance, and fish farms are major sources of resistant *Aeromonas* strains [74–78]. Therefore, there is an urgent need for alternative control strategies.

This study introduces a novel concept as a promising approach for aquaculture: temporarily rearing fish in sediment to introduce ‘wild-type’ gut microbiota. The ‘dirt is good’ hypothesis [79] challenges traditional aquaculture practices that emphasise sterile environments. Although maintaining hygiene remains important, this study shows that introducing beneficial environmental microbes can prevent the overgrowth of opportunistic pathogens, such as *Aeromonas*, which tend to proliferate under domesticated conditions, in the guts of domesticated fish. Previous studies have shown that exposure to diverse microbial communities enhances host immunity in medaka and other species [49, 80, 81], supporting the notion that controlled microbial exposure may be a key strategy for disease prevention. This approach could reduce the dependence on antibiotics and chemical treatments in aquaculture, offering more sustainable disease management practices.

In conclusion, this study provides compelling evidence that ‘wild-type’ gut microbiota can suppress the opportunistic pathogen *Aeromonas* in medaka under domesticated conditions. These findings highlight the crucial role of microbial diversity in maintaining gut health and suggest that introducing environmental microbes could offer a sustainable strategy for disease prevention in aquaculture. Furthermore, by shedding light on the relationship between animals and the environmental microbes in their habitats, this research underscores the significance of understanding these interactions for improving animal health and welfare. While we observed marked differences in microbiome composition, the underlying mechanisms, especially those related to *Aeromonas* suppression in ‘wild-type’ gut microbiota, remain unclear. Further research is needed to identify specific bacterial genera in wild gut microbiomes that inhibit pathogen growth, as well as to assess host responses, including immunity and stress, through survival tests, histological analyses, and immune response evaluations.

## Acknowledgements

We are deeply grateful to Prof. Mitsuru Sakaizumi (Niigata University) for kindly providing some of the wild medaka populations and for guidance on how to catch wild medaka. We are also grateful to Prof. Masakane Yamashita (Hokkaido University) for valuable guidance on maintaining medaka in the laboratory, and to Manabu Murakami (Tokai University) and Yuki Yamada (Ehime University) for invaluable help and advice on sampling in Ehime and Kochi prefectures. We would like to thank Haruka Ooi (AIST) for assistance with experiments and Madoka Miyazaki (AIST) for creating the medaka icons. We also thank Enago (www.enago.jp) for the English language review. This work was financially supported by a Grant-in-Aid for Scientific Research (KAKENHI grant number JP15K21660) from the Japan Society for the Promotion of Science (JSPS), Japan.

## Conflicts of interest

The authors declare no conflicts of interest.

## Author contributions

K.K. and H.I. designed the study with insights from M.K. and Y.K., and H.I. supervised the project. K.K., K.Kb., Y.S., Y.K. and H.I. collected fish and sediment samples. K.K., K.Kb., Y.S., T.H. and H.I. conducted laboratory experiments. K.K. and H.I. analysed and interpreted the data. K.K. and H.I. wrote the paper with substantial input from all authors. All authors reviewed the manuscript.

## Notes

### Competing Interest Statement

The authors have declared no competing interest.

### Summary of Updates

Sections on the background and methods have been updated to clarify motivation and experimental design of this study.

