## Supplementary information for "Wild gut microbiome suppresses the opportunistic pathogen *Aeromonas* in medaka under domesticated rearing conditions"

**Supplementary figures:**

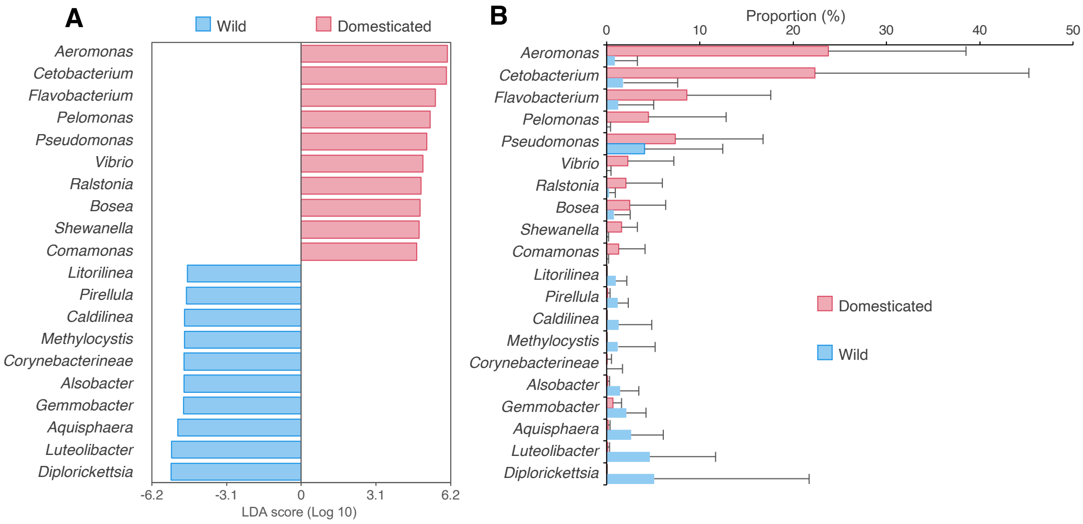


**Fig. S1** Specific bacterial groups in the gut microbiome of domesticated (D) or wild (W) groups. (A) Specific bacterial groups in D or W groups identified via LEfSe analysis. The top 10 genera with the highest LDA scores for D or W groups are shown. (B) Proportion of these specific bacterial groups in the gut microbiomes of each medaka.

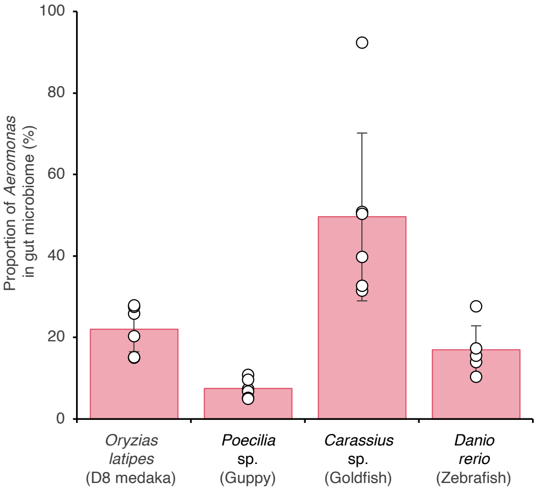


**Fig. S2** Proportion of *Aeromonas* in the gut microbiomes of domesticated fish species. Guppy, goldfish, and zebrafish examined in this experiment were maintained in the same aquarium shop where D8 medaka were purchased.

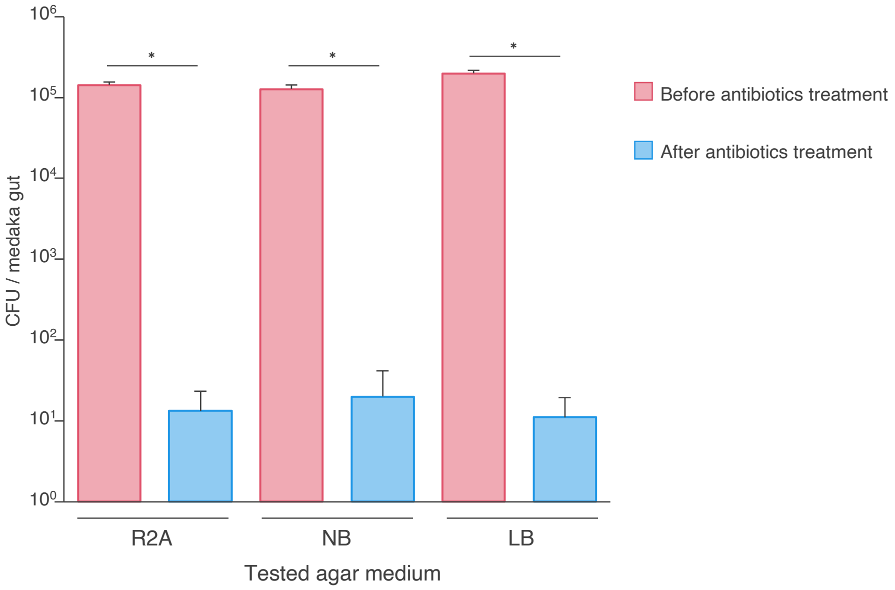


**Fig. S3** Effect of antibiotics treatment on microbial abundance in the medaka gut, measured by colony-forming units (CFUs) on the 1.5% agar plates with three different media: Reasoner's 2A broth (R2A), Nutrient Broth (NB), and Luria-Bertani (LB) broth. Asterisks indicate significant differences ($p < 0.05$, Mann–Whitney U-test).

### Supplementary tables:

Table S1. Identification and haemolytic activity of bacterial strains isolated from medaka gut in the D8 group.

| | | | | Closest reference type strain | | Closest *Aeromonas* strain isolated in this study | |
|---|---|---|---|---|---|---|---|
| Strain name | Haemolytic activity | Sequence length (bp) | Accesession No. of sequence | Reference type strain name | Indetity (%) | Aeromonas strain name | idendity (%) |
| KK93 | + | 1413 | PP474260 | Aeromonas veronii CECT 4257$^T$ | 99.79 | Aeromonas sp. KK125 | 98.58 |
| KK122 | + | 753 | PP549925 | Aeromonas veronii CECT 4257$^T$ | 100 | Aeromonas sp. KK93 | 100 |
| KK123 | + | 753 | PP549926 | Aeromonas veronii CECT 4257$^T$ | 100 | Aeromonas sp. KK93 | 100 |
| KK124 | + | 753 | PP549927 | Aeromonas dhakensis CIP107500$^T$ | 100 | Aeromonas sp. KK125 | 100 |
| KK125 | + | 1412 | PP474261 | Aeromonas dhakensis CIP107500$^T$ | 100 | Aeromonas sp. KK93 | 98.58 |
| KK200 | + | 825 | PP549928 | Pseudomonas alcaligenes NBRC 14159$^T$ | 100 | - | |
| KK203 | + | 753 | PP549924 | Aeromonas veronii CECT 4257$^T$ | 100 | Aeromonas sp. KK93 | 100 |
| KK204 | + | 753 | PP549924 | Aeromonas veronii CECT 4257$^T$ | 100 | Aeromonas sp. KK93 | 100 |
| KK205 | + | 753 | PP549924 | Aeromonas veronii CECT 4257$^T$ | 100 | Aeromonas sp. KK93 | 100 |
| KK206 | + | 753 | PP549924 | Aeromonas veronii CECT 4257$^T$ | 100 | Aeromonas sp. KK93 | 100 |
| KK207 | + | 753 | PP549924 | Aeromonas veronii CECT 4257$^T$ | 100 | Aeromonas sp. KK93 | 100 |
| KK208 | + | 753 | PP549924 | Aeromonas veronii CECT 4257$^T$ | 100 | Aeromonas sp. KK93 | 100 |
| KK209 | + | 753 | PP549924 | Aeromonas veronii CECT 4257$^T$ | 100 | Aeromonas sp. KK93 | 100 |
| KK210 | + | 801 | PP549931 | Pseudomonas alcaligenes NBRC 14159$^T$ | 99.75 | - | |
| KK211 | + | 753 | PP549924 | Aeromonas veronii CECT 4257$^T$ | 100 | Aeromonas sp. KK93 | 100 |
| KK212 | - | 760 | PP549939 | Shewanella xiamenensis S4$^T$ | 98.95 | - | |
| KK213 | - | 798 | PP549952 | Pseudomonas knackmussii | 97.49 | - | |
| KK214 | - | 620 | PP549940 | Shewanella profunda DSM 15900$^T$ | 99.03 | - | |
| KK215 | - | 798 | PP549951 | Pseudomonas alloputida Kh7$^T$ | 100 | - | |
| KK216 | - | 760 | PP549938 | Shewanella xiamenensis S4$^T$ | 99.21 | - | |
| KK219 | - | 711 | PP549937 | Pseudomonas furukawaii KF707$^T$ | 100 | - | |
| KK220 | - | 783 | PP549961 | Fictibacillus phosphorivorans | 100 | - | |
| KK221 | - | 718 | PP549960 | Pseudomonas knackmussii B13$^T$ | 97.91 | - | |
| KK223 | - | 760 | PP549941 | Shewanella profunda DSM 15900$^T$ | 99.21 | - | |
| KK224 | + | 825 | PP549929 | Pseudomonas alcaligenes NBRC 14159$^T$ | 100 | - | |
| KK225 | + | 825 | PP549930 | Pseudomonas alcaligenes NBRC 14159$^T$ | 100 | - | |
| KK227 | + | 753 | PP549924 | Aeromonas veronii CECT 4257$^T$ | 100 | Aeromonas sp. KK93 | 100 |
| KK228 | + | 753 | PP549924 | Aeromonas veronii CECT 4257$^T$ | 100 | Aeromonas sp. KK93 | 100 |
| KK229 | + | 753 | PP549924 | Aeromonas veronii CECT 4257$^T$ | 100 | Aeromonas sp. KK93 | 100 |
| KK230 | + | 753 | PP549924 | Aeromonas veronii CECT 4257$^T$ | 100 | Aeromonas sp. KK93 | 100 |
| KK231 | + | 753 | PP549924 | Aeromonas veronii CECT 4257$^T$ | 100 | Aeromonas sp. KK93 | 100 |
| KK232 | + | 753 | PP549924 | Aeromonas veronii CECT 4257$^T$ | 100 | Aeromonas sp. KK93 | 100 |
| KK233 | + | 753 | PP549924 | Aeromonas veronii CECT 4257$^T$ | 100 | Aeromonas sp. KK93 | 100 |
| KK235 | + | 753 | PP549924 | Aeromonas veronii CECT 4257$^T$ | 100 | Aeromonas sp. KK93 | 100 |
| KK236 | - | 799 | PP549950 | Pseudomonas alloputida Kh7$^T$ | 100 | - | |
| KK237 | - | 753 | PP549949 | Pseudomonas knackmussii B13$^T$ | 98.01 | - | |
| KK238 | - | 744 | PP549959 | Ensifer morelensis Lc04$^T$ | 99.32 | - | |
| KK239 | - | 760 | PP549942 | Shewanella profunda DSM 15900$^T$ | 99.21 | - | |
| KK248 | + | 753 | PP549924 | Aeromonas veronii CECT 4257$^T$ | 100 | Aeromonas sp. KK93 | 100 |
| KK249 | + | 753 | PP549924 | Aeromonas veronii CECT 4257$^T$ | 100 | Aeromonas sp. KK93 | 100 |
| KK250 | + | 753 | PP549924 | Aeromonas veronii CECT 4257$^T$ | 100 | Aeromonas sp. KK93 | 100 |
| KK251 | + | 753 | PP549924 | Aeromonas veronii CECT 4257$^T$ | 100 | Aeromonas sp. KK93 | 100 |
| KK252 | + | 687 | PP549935 | Shewanella putrefaciens JCM 20190$^T$ | 99.26 | - | |
| KK253 | + | 750 | PP549936 | Crenobacter intestini GY 70310$^T$ | 99.60 | - | |
| KK255 | + | 825 | PP549934 | Pseudomonas alloputida Kh7$^T$ | 100 | - | |
| KK256 | + | 825 | PP549933 | Pseudomonas alcaligenes NBRC 14159$^T$ | 100 | - | |
| KK260 | - | 762 | PP549944 | Aeromonas veronii CECT 4257$^T$ | 100 | Aeromonas sp. KK93 | 100 |
| KK261 | - | 762 | PP549945 | Aeromonas veronii CECT 4257$^T$ | 100 | Aeromonas sp. KK93 | 100 |
| KK262 | - | 762 | PP549946 | Aeromonas veronii CECT 4257$^T$ | 100 | Aeromonas sp. KK93 | 100 |
| KK263 | - | 762 | PP549947 | Aeromonas veronii CECT 4257$^T$ | 100 | Aeromonas sp. KK93 | 100 |
| KK264 | - | 702 | PP549963 | Flavobacterium cutihirudinis DSM 25795$^T$ | 99.42 | - | |
| KK268 | - | 753 | PP549948 | Pseudomonas knackmussii B13$^T$ | 98.01 | - | |
| KK270 | - | 770 | PP549964 | Microbacterium aureliae JF-6$^T$ | 98.44 | - | |
| KK271 | - | 760 | PP549943 | Shewanella profunda DSM 15900$^T$ | 99.21 | - | |
| KK272 | + | 753 | PP549924 | Aeromonas veronii CECT 4257$^T$ | 100 | Aeromonas sp. KK93 | 100 |
| KK273 | + | 753 | PP549924 | Aeromonas veronii CECT 4257$^T$ | 100 | Aeromonas sp. KK93 | 100 |
| KK274 | + | 753 | PP549924 | Aeromonas veronii CECT 4257$^T$ | 100 | Aeromonas sp. KK93 | 100 |
| KK275 | + | 753 | PP549924 | Aeromonas veronii CECT 4257$^T$ | 100 | Aeromonas sp. KK93 | 100 |
| KK276 | + | 753 | PP549924 | Aeromonas veronii CECT 4257$^T$ | 100 | Aeromonas sp. KK93 | 100 |
| KK277 | + | 753 | PP549924 | Aeromonas veronii CECT 4257$^T$ | 100 | Aeromonas sp. KK93 | 100 |
| KK278 | + | 825 | PP549932 | Pseudomonas alcaligenes NBRC 14159$^T$ | 99.78 | - | |
| KK279 | - | 720 | PP549953 | Edwardsiella ictaluri ATCC33202$^T$ | 99.86 | - | |
| KK280 | - | 720 | PP549954 | Edwardsiella ictaluri ATCC33202$^T$ | 99.86 | - | |
| KK281 | - | 720 | PP549955 | Edwardsiella ictaluri ATCC33202$^T$ | 99.86 | - | |
| KK282 | - | 720 | PP549956 | Edwardsiella ictaluri ATCC33202$^T$ | 99.86 | - | |
| KK283 | - | 720 | PP549957 | Edwardsiella ictaluri ATCC33202$^T$ | 99.86 | - | |
| KK284 | - | 720 | PP549958 | Edwardsiella ictaluri ATCC33202$^T$ | 99.86 | - | |
| KK285 | - | 778 | PP549962 | Chryseobacterium cucumeris GSE06$^T$ | 100 | - | |

Table S2. Dominant members of the gut microbiome in medaka reared for one week with sediment from the site where W6 group were collected.

| Taxonomy name[a] | Gut[b] | Sediment[c] | Water[d] |
|---|---|---|---|
| *Aquisphaera* | 13.28 ± 0.84 | 0.38 ± 0.02 | 0.07 ± 0.02 |
| *Pirellula* | 4.72 ± 1.84 | 0.70 ± 0.11 | 0.01 ± 0.01 |
| Verrucomicrobia Subdivision3 | 4.28 ± 0.71 | 2.45 ± 0.20 | 0.07 ± 0.02 |
| *Reyranella* | 3.70 ± 0.87 | 0.04 ± 0.003 | 0.03 ± 0.007 |
| *Zavarzinella* | 3.18 ± 0.66 | 0.20 ± 0.03 | 0.005 ± 0.007 |
| *Singulisphaera* | 2.14 ± 0.26 | 0.17 ± 0.02 | 0.002 ± 0.004 |
| *Acidobacteria* Gp16 | 2.13 ± 0.30 | 0.90 ± 0.07 | 0.009 ± 0.008 |
| *Gemmata* | 2.03 ± 0.45 | 0.10 ± 0.03 | 0 |
| *Litorilinea* | 1.80 ± 0.17 | 0.82 ± 0.07 | 0 |
| *Thermogutta* | 1.70 ± 0.15 | 0.54 ± 0.04 | 0.008 ± 0.01 |
| *Anaeromyxobacter* | 1.68 ± 0.11 | 1.75 ± 0.13 | 0.32 ± 0.05 |
| *Alsobacter* | 1.65 ± 0.10 | 0.13 ± 0.01 | 0.38 ± 0.002 |
| *Geobacter* | 1.39 ± 0.84 | 4.57 ± 0.10 | 2.37 ± 0.15 |
| *Simkania* | 1.35 ± 0.53 | 0.01 ± 0.01 | 0 |
| *Desulfovibrio* | 1.27 ± 0.57 | 0.05 ± 0.04 | 0.005 ± 0.003 |
| *Spartobacteria* | 1.20 ± 0.08 | 0.16 ± 0.03 | 0.009 ± 0.007 |
| *Rubinisphaera* | 1.08 ± 0.43 | 0.03 ± 0.03 | 0.008 ± 0.006 |
| *Gemmobacter* | 1.03 ± 0.13 | 3.91 ± 0.31 | 1.45 ± 0.10 |

[a]Only members with proportion >1.0% in medaka gut microbiome are listed.

[b]Proportion (%) in medaka gut microbiome.

[c]Proportion (%) in sediment microbiome.

[d]Proportion (%) in water microbiome.

Table S3. Dominant members of the gut microbiome in medaka reared for one week with sediment from the site where W12 group were collected.

| Taxonomy name[a] | Gut[b] | Sediment[c] | Water[d] |
|---|---|---|---|
| *Zavarzinella* | 11.85 ± 1.58 | 0.53 ± 0.03 | 0.01 ± 0.01 |
| *Pirellula* | 8.62 ± 3.94 | 0.65 ± 0.07 | 0.02 ± 0.001 |
| *Aquisphaera* | 8.49 ± 0.79 | 0.19 ± 0.08 | 1.72 ± 0.18 |
| *Reyranella* | 5.69 ± 1.69 | 0.11 ± 0.04 | 0.09 ± 0.008 |
| *Aeromonas* | 2.60 ± 1.72 | 0.01 ± 0.01 | 0.02 ± 0.005 |
| *Gemmobacter* | 2.36 ± 1.18 | 14.46 ± 0.47 | 28.27 ± 0.46 |
| *Luteolibacter* | 2.29 ± 0.85 | 1.12 ± 0.08 | 0.007 ± 0.002 |
| Verrucomicrobia Subdivision3 | 2.06 ± 0.45 | 2.37 ± 0.12 | 0.009 ± 0.005 |
| *Thermogutta* | 1.89 ± 0.18 | 0.67 ± 0.13 | 0.004 ± 0.003 |
| *Litorilinea* | 1.82 ± 0.67 | 0.29 ± 0.03 | 0 |
| *Bdellovibrio* | 1.29 ± 1.26 | 0.19 ± 0.02 | 0.11 ± 0.003 |
| *Gemmata* | 1.02 ± 0.14 | 0.25 ± 0.04 | 0 |

[a]Only members with proportion >1.0% in medaka gut microbiome are listed.

[b]Proportion (%) in medaka gut microbiome.

[c]Proportion (%) in sediment microbiome.

[d]Proportion (%) in water microbiome.

Table S4. Dominant members of the gut microbiome in medaka reared for one week with sediment from the site where W14 group were collected.

| Taxonomy name[a] | Gut[b] | Sediment[c] | Water[d] |
|---|---|---|---|
| *Aquisphaera* | 14.95 ± 2.70 | 0.72 ± 0.21 | 3.56 ± 0.16 |
| *Zavarzinella* | 10.49 ± 0.55 | 0.29 ± 0.04 | 0.03 ± 0.005 |
| *Pirellula* | 9.03 ± 1.11 | 1.52 ± 0.17 | 1.89 ± 0.21 |
| *Reyranella* | 4.96 ± 0.91 | 0.06 ± 0.01 | 0.22 ± 0.02 |
| *Cetobacterium* | 2.89 ± 1.57 | 0.44 ± 0.13 | 0.24 ± 0.006 |
| *Acidobacteria* Gp16 | 2.65 ± 1.13 | 1.76 ± 0.08 | 0.002 ± 0.003 |
| *Gemmata* | 2.59 ± 0.15 | 0.13 ± 0.05 | 0.003 ± 0.004 |
| *Nitrolancea* | 2.28 ± 0.43 | 0.02 ± 0.02 | 0.003 ± 0.004 |
| *Spartobacteria* | 2.06 ± 0.92 | 0.14 ± 0.03 | 0.002 ± 0.003 |
| *Singulisphaera* | 1.86 ± 0.28 | 0.16 ± 0.03 | 0 |
| *Aeromonas* | 1.66 ± 0.35 | 0.39 ± 0.05 | 0.11 ± 0.01 |
| *Acidobacteria* Gp6 | 1.58 ± 0.48 | 0.91 ± 0.20 | 0.002 ± 0.003 |
| *Litorilinea* | 1.56 ± 0.42 | 1.87 ± 0.26 | 0.002 ± 0.003 |
| *Thermogutta* | 1.40 ± 0.13 | 0.34 ± 0.06 | 0.002 ± 0.003 |
| Verrucomicrobia Subdivision3 | 1.24 ± 0.25 | 1.79 ± 0.07 | 0.12 ± 0.06 |
| *Caldilinea* | 1.16 ± 0.20 | 0.51 ± 0.03 | 0 |
| *Thermomarinilinea* | 1.14 ± 0.56 | 1.43 ± 0.12 | 0 |

[a]Only members with proportion >1.0% in medaka gut microbiome are listed.

[b]Proportion (%) in medaka gut microbiome.

[c]Proportion (%) in sediment microbiome.

[d]Proportion (%) in water microbiome.

Table S5. Dominant members of the gut microbiome in medaka reared for one week with sediment from the site where W15 group were collected.

| Taxonomy name[a] | Gut[b] | Sediment[c] | Water[d] |
|---|---|---|---|
| *Aquisphaera* | 16.95 ± 1.96 | 0.59 ± 0.09 | 0.01 ± 0.004 |
| *Reyranella* | 7.81 ± 2.10 | 0.30 ± 0.08 | 0.07 ± 0.01 |
| *Pirellula* | 7.43 ± 1.40 | 0.34 ± 0.21 | 0.01 ± 0.004 |
| *Gemmobacter* | 3.62 ± 2.63 | 0.39 ± 0.06 | 8.67 ± 0.25 |
| *Zavarzinella* | 3.43 ± 0.67 | 0.19 ± 0.13 | 0.005 ± 0.004 |
| *Luteolibacter* | 1.43 ± 0.18 | 0.16 ± 0.09 | 0.07 ± 0.03 |
| *Zoogloea* | 1.32 ± 1.80 | 0.38 ± 0.05 | 0.71 ± 0.04 |
| Verrucomicrobia Subdivision3 | 1.32 ± 0.62 | 1.66 ± 0.12 | 0.02 ± 0.01 |
| *Alsobacter* | 1.29 ± 0.05 | 0.26 ± 0.10 | 0.03 ± 0.01 |
| *Singulisphaera* | 1.18 ± 0.13 | 0.18 ± 0.03 | 0.01 ± 0.004 |
| *Acidobacteria* Gp16 | 1.06 ± 0.69 | 2.26 ± 0.13 | 0 |
| *Rhodopirellula* | 1.04 ± 0.47 | 0 | 0 |

[a]Only members with proportion >1.0% in medaka gut microbiome are listed.

[b]Proportion (%) in medaka gut microbiome.

[c]Proportion (%) in sediment microbiome.

[d]Proportion (%) in water microbiome.

Table S6. Dominant bacterial members of the rearing water used for Step 3 in Fig. 5A.

| Taxonomy name[a] | Water[b] |
|---|---|
| *Cetobacterium* | 6.82 ± 3.84 |
| *Verrucomicrobium* | 5.81 ± 3.95 |
| *Sphingobium* | 11.29 ± 1.55 |
| *Aquabacterium* | 8.97 ± 1.48 |
| *Acidovorax* | 8.87 ± 1.63 |
| *Aeromonas* | 2.17 ± 1.51 |
| *Planctopirus* | 1.88 ± 1.53 |
| *Nevskia* | 4.76 ± 1.00 |
| *Flavobacterium* | 1.68 ± 1.04 |
| *Reyranella* | 1.88 ± 1.53 |
| *Comamonas* | 1.56 ± 0.19 |
| *Flectobacillus* | 1.74 ± 0.50 |
| *Elstera* | 1.37 ± 0.30 |
| *Azotobacter* | 1.05 ± 0.30 |

[a]Only members with proportion >1.0% in rearing water are listed.

[b]Proportion (%) in water microbiome.
